# Comparative analysis of NSP5/VP2-induced viroplasm-like structures in rotavirus species A to J

**DOI:** 10.1101/2025.06.09.658619

**Authors:** Ariana Cosic, Melissa Lee, Kurt Tobler, Claudio Aguilar, Cornel Fraefel, Catherine Eichwald

## Abstract

Rotavirus (RV) is classified into nine species, A-D and F-J, with RV species A (RVA) being the most extensively studied. While RVA infects infants and young animals, non-RVA species infect adult humans, various mammals, and birds. However, the lack of appropriate research tools has limited our understanding of non-RVA life cycles. RVA replication and assembly occur in cytosolic inclusions termed viroplasms. We recently identified viroplasm-like structures (VLS) composed of NSP5 and NSP2, designated as VLS(NSP2)i, in non-RVA. In this context, globular VLS(NSP2)i formed in RVA, RVB, RVD, RVF, RVG, and RVI, but not in RVC, RVH, and RVJ. Additionally, in RVA, VLS can also be formed through co-expression of NSP5 with VP2, referred to as VLS(VP2)i. Here, we report VLS(VP2)i formation in all non-RVA species except RVB, with notable VLS formation in RVH and RVJ. Moreover, NSP2 RVH or RVJ are recruited into VLS(VP2)i. The NSP5 C-terminal region in non-RVA is required for association with VP2 and forming VLS(VP2)i. Mutation of conserved VP2-L141 in RVA to alanine disrupts viroplasm formation, impairing RV replication. Equivalent residues within the same predicted VP2 region disrupt VLS formation across non-RVA. We also observed interspecies VLS formation, particularly between closely related RVA and RVC, RVH and RVJ, and RVD and RVF. Interestingly, substituting the N-terminal region of VP2 RVB with that of the closely related VP2 RVG restores its ability to form VLS with NSP5 RVB. Elucidating the formation of viroplasms is essential for developing strategies to halt infection across RV species A to J.

**Importance:** Rotaviruses (RV) are a group of viruses classified into species A through J, with species A being the best understood. Other RV species infecting animals and humans are less studied due to limited research tools. In RVA, the virus replicates in specialized compartments called viroplasms formed in the cytoplasm by viral proteins, including NSP5, NSP2, and VP2. In this study, we explored how similar structures, termed viroplasm-like structures (VLS), are formed by proteins of other RV species. We found that in most species, NSP5 and VP2 form VLSs. One exception was RVB, where VLS formation was observed only when the VP2 protein was mutated to resemble that of a related species. We also identified key regions in the VP2 protein that are essential for forming these structures. Understanding how viroplasms form across different RV species may help develop new strategies to block infection in humans and animals.

## Introduction

Rotavirus (RV) is an etiological agent belonging to the *Sedoreoviridae* family and is responsible for severe gastroenteritis and dehydration. According to the International Committee on the Taxonomy of Viruses (ICTV), RVs are currently grouped into nine species, from *Rotavirus alphagastroenteritidis* to *Rotavirus jotagastroenteritidis* (1). For simplicity, the RV species will be termed herewith as A to D and F to J. The RV species A (RVA) is the prevailing RV species among infants and young children, killing approximately 128,000 children per year, mainly in low- and middle-income countries (2). RVA also has a broad spectrum of strains primarily infecting young mammals like piglets and calves. The non-RVA species have been isolated from diverse hosts, including mammals and avians. Outbreaks from RVB, RVC, and RVH are the leading cause of diarrhea among the adult human population in several countries (3–7). In the US, RV infections are the second most common cause of diarrhea in adults after norovirus (8–10). From a veterinary perspective, RV infections significantly impact livestock worldwide. RV accounts for 80% of diarrhea cases in piglets in the USA, Canada, and Mexico, with potential zoonotic implications in humans (11). RV species D, F, and G have only been detected in avian species, affecting the poultry industry by impacting the feed conversion ratio and resulting in substantial economic losses (12). All the information compiled on the RV replication cycle is based on RVA. Studying the replication of non-RVA species is challenging, and as a result, their biology remains poorly understood. The few isolated viruses of non-RVA species do not replicate in tissue culture (13), and tools recognizing their specific proteins, like specific antibodies, are unavailable.

During RVA infection, the external layer is lost after virion internalization, and transcriptionally active double-layered particles (DLP) are released into the cytosol (14). The newly released transcripts initiate the synthesis of viral proteins necessary for viral replication. Among those proteins, the nonstructural proteins NSP2 and NSP5 and the structural proteins VP1, VP2, VP3, and VP6 comprise part of the RV viral factories termed viroplasms (15). The viroplasms correspond to electron-dense membrane-less globular cytosolic inclusions where viral genome transcription, replication, and the packaging of the newly synthesized pre-genomic RNA segments into the viral cores occur. The viroplasms are highly dynamic, being able to coalesce between them and migrate to the juxtanuclear region of the cell at later stages post-infection (16–18). Furthermore, despite not yet being well-defined, several host factors have been identified as necessary for viroplasm formation and maintenance (19–22). For RVA, the initiation process for viroplasm formation requires a scaffold of lipid droplets that incorporates perilipin-1 (23, 24). Furthermore, the host cytoskeleton, actin filaments, and microtubules (MT) play a role in the formation, maintenance, and dynamics of the viroplasms (17, 25, 26). In this context, NSP2 octamers are directly associated with MTs to promote viroplasm coalescence (17, 27–30). Moreover, VP2 plays a role in viroplasm dynamics by allowing its perinuclear motion (17). Consistent with these features, the viroplasms are considered liquid-liquid phase-separated structures (31). Interestingly, the co-expression of NSP5 with either NSP2 or VP2 leads to the formation of cytosolic inclusions named viroplasm-like structures (VLS), which are morphologically similar to viroplasms but unable to yield viral progeny (16, 17, 32–35). When associating with NSP2 or VP2, NSP5 is primed at serine-67 by the casein kinase 1 alpha, triggering NSP5 hyperphosphorylation (32, 36–38). The NSP5 S67A mutation prevents viroplasm formation (39). The NSP5 phosphorylation is consistent with a trait for recently described liquid-liquid phase separation conditions of the viroplasms (31). NSP5 is not only required for viroplasm formation and virus replication (39–41) but also plays a multifunctional role in the RV life cycle, interacting with NSP6 (35), NSP2 (16), VP1 (42), VP2 (43, 44), and unspecifically with dsRNA (45). These attributes are consistent with its predicted disordered nature (46–48). Interestingly, the C-terminal ordered region (henceforth tail) of NSP5 is needed for its self-oligomerization (35, 36), to associate with other RV proteins (16, 35, 42, 44), and to form the viroplasms (39). NSP5 is sumoylated (49), presumably a pre-requirement for interacting with viral or host components. Collectively, NSP5 is a crucial component in RV replication.

RVA NSP2 self-assembles into an octamer composed of two doughnut-shaped tetramers, exhibiting a highly electropositive surface (28, 50). NSP2 is associated with several enzymatic functions, including nucleoside diphosphate kinase-like activity (51), RNA-helix destabilization (51), and nucleoside triphosphatase activity (28), all of which are consistent with molecular motor properties. Moreover, NSP2 phosphorylation has been linked to viroplasm formation, as shown by the delayed appearance of viroplasms in cells infected with a recombinant RV expressing a phosphomimetic mutant NSP2 S313D (52). NSP2 directly contributes to viroplasm coalescence (16), a process involving its association with MTs (29). Additionally, the flexible C-terminal region of NSP2 enhances viroplasm morphology (53) and RNA chaperone activity (27). Notably, NSP2 binds both to VP1 and viral RNA (54, 55), implicating it as a key component of replication intermediates within the viroplasms.

Likewise, the core-shell protein VP2, in addition to its structural role in safeguarding the RVA genome, can activate and regulate the RNA-dependent RNA polymerase (RdRp) VP1, allowing for genome replication. VP2 forms asymmetric decameric structures converging in the five-fold axis, which cannot be dissociated (48, 56–59). Each decameric subunit comprises a main domain of VP2 (residues ∼ 100 to 880), creating a thin, comma-shaped plate where the unfolded N-terminal domain (NTD) is positioned beneath the decameric five-fold axis. Several viral proteins (58, 60–62) and nonspecific ssRNA (63) interact with VP2, primarily to facilitate association with the NTD. These interactions are closely linked to the core-shell structure and genome replication. Additionally, VP2 serves as a key component in forming viroplasms and, when co-expressed with NSP5, produces VLS (17, 32, 44, 64). In this context, the VLSs induced by VP2 are dynamic as they migrate to the perinuclear region (17). Furthermore, the highly conserved L124 of VP2 in RVA is crucial for its association with NSP5. When L124 is mutated to alanine, VP2 L124A disrupts viroplasm morphology, rendering RV replication incompetent (44). Recently, it has been suggested that VP2 may have further roles early post-infection due to its interaction with NSP2, which prevents its spontaneous oligomerization and sumoylation, thereby enhancing the ability of VP2 to interact with other proteins (17, 49).

We recently investigated the ability of NSP5 and NSP2 from non-RVA species to form VLSs (65).The co-expression of these two proteins resulted in the formation of globular VLSs in RVA, RVB, RVD, RVF, RVG, and RVI, while RVC formed filamentous VLSs. In contrast, the co-expression of NSP5 and NSP2 from RV species H and J did not lead to VLS formation. Interestingly, NSP5 from all RV species oligomerizes through its tail region. Except for NSP5 from RVJ, all NSP5 orthologs interacted with their respective NSP2 proteins. We also found that interspecies VLSs formed between closely related RV species, specifically B and G, and between D and F. Furthermore, VLSs were observed in RVH and RVJ when the tail of NSP5-RVH or NSP5-RVJ was substituted with that of NSP5 from RVA and co-expressed with the respective NSP2.

In this study, we characterized the formation of VLS supported by the co-expression of NSP5 and VP2 across RV species A to J. We determined that the NSP5 tail is crucial for both VLS formation and its interaction with VP2 in all RV species tested. A point mutation to alanine of a conserved amino acid residue in VP2 disrupts VLS formation. Heterologous VLS formation was observed between closely related RV species pairs: A and C, D and F, as well as H and J. Additionally, we demonstrated that the unstructured N-terminal region of VP2 is necessary for VLS formation.

## Results

### Biophysical features of VP2 in RV species A to J

We recently demonstrated that exploring the mechanism of replication of non-RVA species is possible by extrapolating the role of NSP5 and NSP2 from RVA to their orthologs in other RV species (65). In this context, RVA also forms VLS upon co-expression of NSP5 with VP2 (32, 44, 66). Therefore, we wondered whether VP2 of non-RVA could have a similar role in VLS formation when co-expressed with its cognate NSP5. First, we identified the available open reading frames of VP2 for RV species A to J in the NCBI database that form pairs with their cognate NSP5 and NSP2 described previously (65). However, this option was not feasible for VP2 of RVB, RVC, and RVI, since the complete sequences were unavailable and were therefore replaced by other strains with higher homology (**Table 1**)(65). The VP2 of diverse RV species differs in their amino acid length (**Table 1 and Fig. 1e**), with variations of up to 109 residues, where the shortest one corresponds to RVA with 882 amino acids and the longest to RVG with 991 amino acids. Moreover, VP2 primary sequences across species A to J are highly diverse compared to our model VP2 RVA corresponding to VP2 simian strain SA11, where the closest related is RVF with 68.51% similarity and the most diverse is RVJ with a 35.28% similarity (**Table 1**). Consistent with the fact that the N-terminal domain of VP2 RVA is unfolded (residues ∼ 1 to 100 for type A and ∼1 to 80 for type B) (57–59, 67), the PONDR score also indicates a N-terminal region predicted to be highly disordered when comparing the primary structures of VP2 RVA of two model strains, simian strain SA11 and porcine strain OSU (**Fig. 1a**). Similarly, VP2 of non-RVA species presents a disordered N-terminal region (**Fig. 1 b and d**), except for VP2 of RVB that lacks this predicted disordered region (**Fig. 1c**). Notably, the predicted disordered N-terminal region of VP2 across diverse RV species can be subdivided according to their N-terminal: completely disordered for RVA, RVC, RVD and RVF (**Fig. 1b**), and partially disordered for RVG, RVH, RVI and RVJ (**Fig. 1d**). The latter have a few ordered residues at their N-termini followed by a disordered region of approximately 50 residues. We compared the predicted folding of VP2 across different RV species and found that the AlphaFold3 prediction of the dimeric structure of VP2 RVA (**Fig. S1**, RVA) closely matched the previously experimental structure (58). For VP2 of non-RVA species, AlphaFold3 predicted similar overall structures, particularly in the apical, central, and dimerization regions (**Fig. S1**). Consistent with the predicted disordered nature of the N-terminal regions of VP2, AlphaFold3 showed reduced confidence for this region across all analyzed RV species (data not shown). Accordingly, we designed a series of plasmids encoding the open reading frame of VP2 of RV species A to J with a Flag tag fused at their N-terminal region (**Fig. 1e**) (44). Lysates of MA104 cells expressing Flag-VP2 of RV species A to J were examined by immunoblotting using a polyclonal anti-VP2 designed for VP2/RVA strain SA11(42) (**Fig. 1f, upper panel**), resulting in the detection of VP2 of RVA and, with less affinity, VP2 of RVB and RVC, suggesting antigenic homology for VP2 among these three RV species but not with other RV species. As expected, the subsequent incubation of the same membrane with an anti-Flag antibody allowed the recognition of all tested RV species of VP2, which proved to migrate at their predicted molecular weight (**Table S1**).

**Figure 1.**
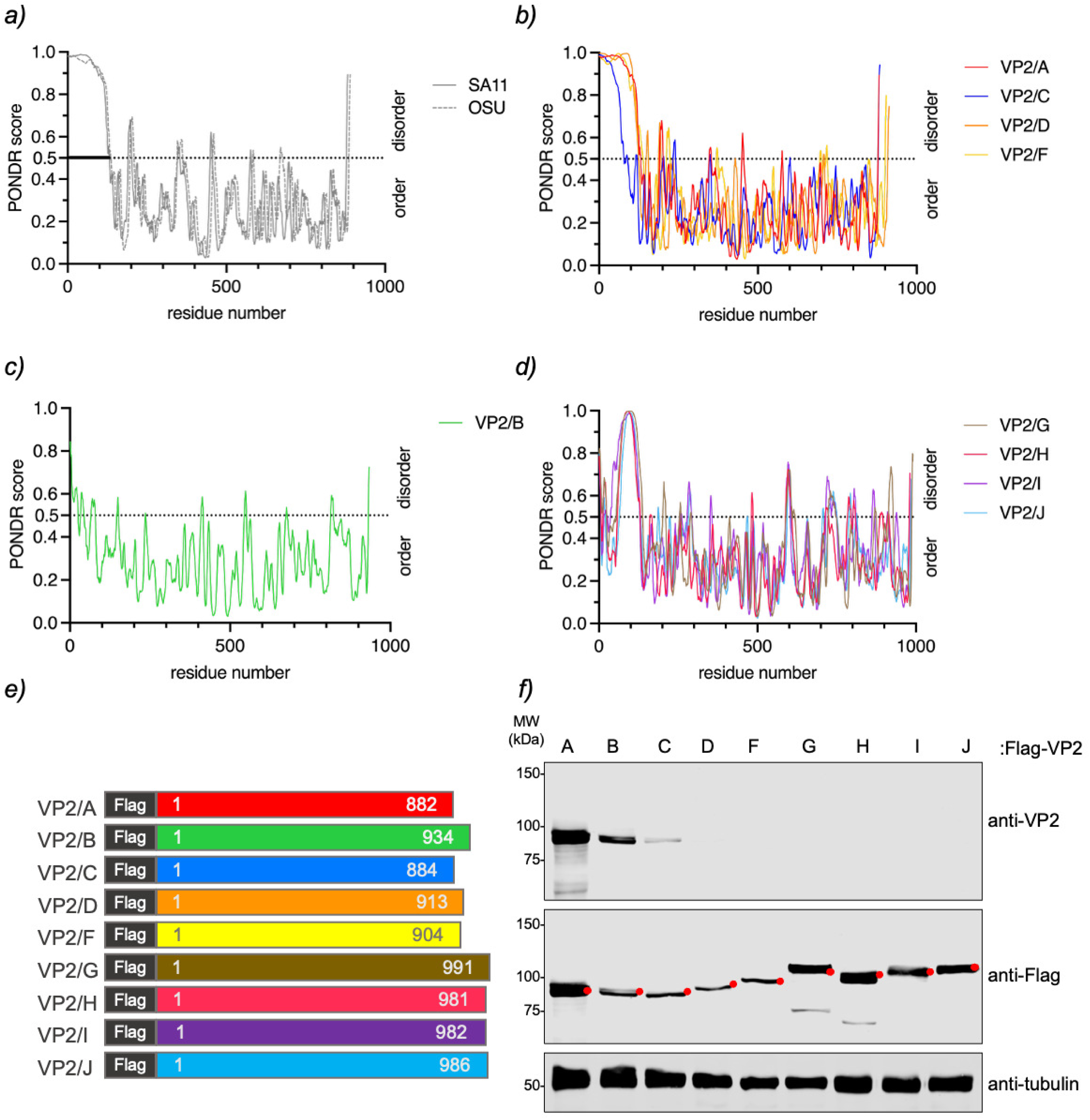
IDR and expression of VP2 in species A to J. Plots comparing the IDR prediction of VP2 from RVA strains OSU and SA11 **(a)**, RVA (strain SA11), RVC, RVD, and RVF **(b)**, RVB **(c)**, and RVG-RVJ **(d)**. **e)** Schematic representation of VP2 from RV species A to J, fused to a Flag tag at its N-terminal region. **f)** Immunoblotting of MA104 cell extracts expressing Flag-VP2 from RV species A to J. The membrane was incubated with guinea pig anti-VP2 (top panel) and mouse mAb anti-Flag (middle panel). Anti-tubulin served as a loading control (bottom panel). The red dot indicates the predicted molecular weight of the recombinant proteins.

**Table 1:**
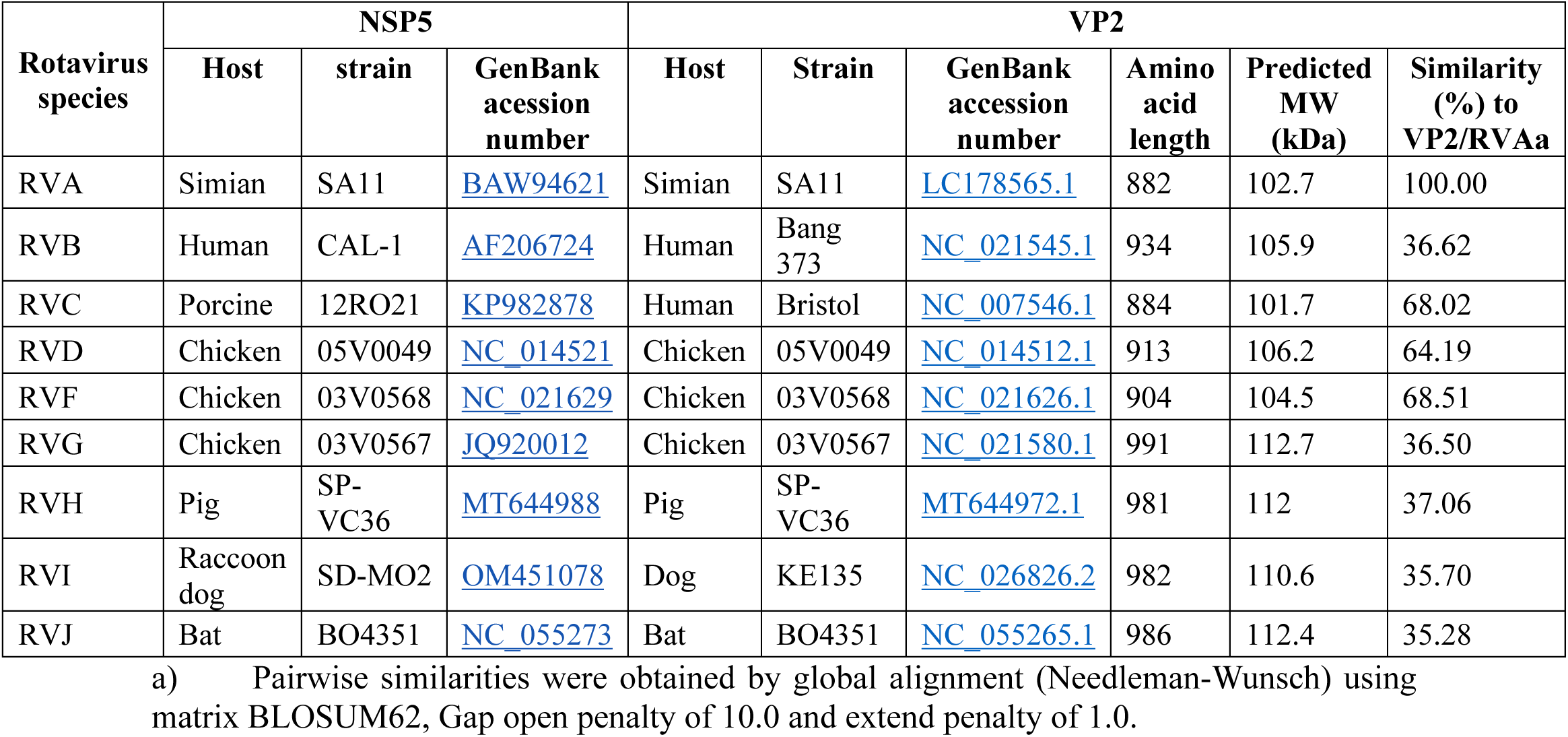
NSP5 and VP2 protein features of the RV species analyzed in this study.

### VLS formation by co-expression of NSP5 with VP2 across RV species A to J

Next, we investigated whether <u>b</u>iotin <u>a</u>cceptor <u>p</u>eptide (BAP)-tagged NSP5(65) co-expressed with their cognate Flag-VP2 protein supports the formation of VLS in RV species A to J. Of note, VLSs are visualized by colocalization of the signals of NSP5 with NSP2 or VP2 in globular cytosolic inclusions (25, 32, 44, 65, 66, 68). In the first instance (**Fig. 2**), the proteins were expressed in mammalian MA/cytBirA cells fixed at 16 h post-transfection (hpt). VLS formation was monitored by immunofluorescence for the detection of NSP5 fused to BAP tag (streptavidin, green) and Flag-VP2 (mAb anti-Flag, red), respectively. As expected (44), NSP5-BAP and Flag-VP2 of RVA colocalized, forming globular cytosolic inclusions corresponding to VLS. Similarly, the co-expression of NSP5-BAP and Flag-VP2 of RVC, RVF, RVG, RVH, and RVJ also led to the formation of globular VLSs. However, the co-expression of these proteins in RVB and RVD did not allow VLS formation. Even though several transfection ratios (4:1, 2:1, 1:1, 1:2, and 1:4) of NSP5-BAP and Flag-VP2 (**Fig. S2a**) were tested for RVB species, none allowed VLS formation. As described previously (65), BAP-NSP5 of RVD and RVF formed globular inclusions in the nuclei.

**Figure 2.**
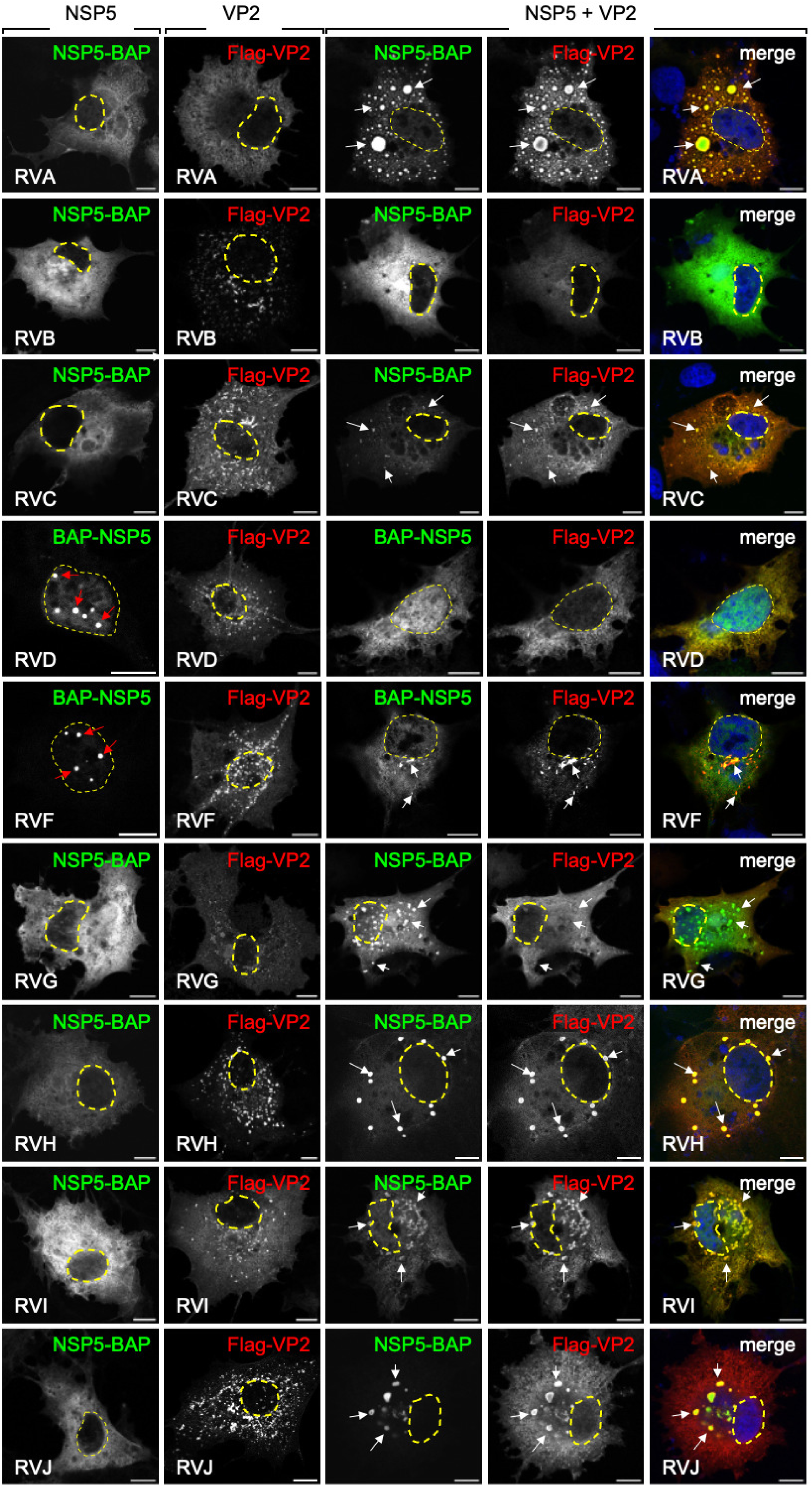
Characterization of VLS formation through co-expression of NSP5-BAP and Flag-VP2 across RV species A to J. Immunofluorescence images of MA/cytBirA cells expressing NSP5-BAP (RVA-RVC and RVG-RVJ) or BAP-NSP5 (RVD and RVF) alone (first column), Flag-VP2 (RVA-RVJ) alone (second column), and the co-expression of both proteins (third, fourth, and fifth columns). At 16 hpt, the cells were fixed and stained to detect NSP5 with Streptavidin (green) and VP2 (anti-Flag, red). A merged image is shown in the fifth column. Nuclei were stained with DAPI (blue). The scale bar is 10 µm. The white and red arrows point to globular VLS and nuclear inclusions, respectively. The yellow discontinuous line marks the nucleus as determined by DAPI staining.

Since RVD, RVF, and RVG were isolated from avian hosts, we wondered whether the host environment could influence the folding and interaction behavior of the corresponding NSP5 and VP2 and, thus, VLS formation. To test this hypothesis, we expressed V5-tagged NSP5 and Flag-VP2 in LMH chicken epithelial cells and monitored VLS formation via immunofluorescence. In this context (**Fig. 3a**), the expression of Flag-VP2 alone from RVD, RVF, and RVG forms filamentous structures for RVD and RVG and globular ones for RVF, contrasting with the diffuse cytosolic aggregates found in MA/cytBirA cells for these RV species. As expected (65), V5-NSP5 of RVD and RVF was diffused in the cytosol, while NSP5-V5 of RVG formed globular cytosolic inclusions (**Fig. 3b)**. Furthermore, upon deletion of the tail region NSP5 of RVD and RVF localized in both the cytosol and the nucleus. As depicted in **Fig. 3c**, the co-expression of NSP5 fused to V5 with Flag-VP2 of RV species D, F, and G leads to the formation of cytosolic globular VLS in LMH cells.

**Figure 3.**
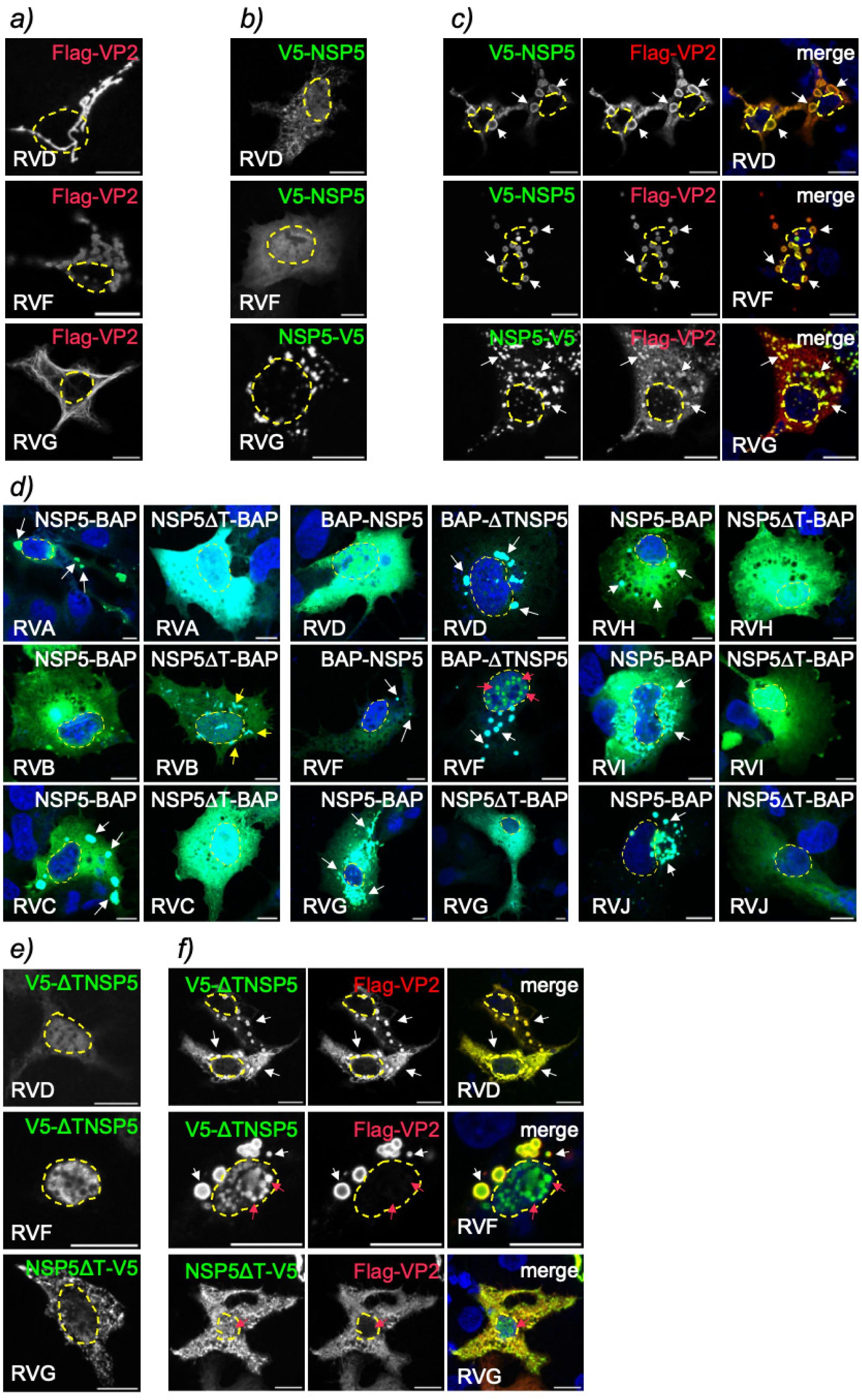
The role of NSP5 tail in the formation of VLS induced by VP2 in RV species A to J. Immunofluorescence images of chicken LMH cells expressing **(a)** Flag-VP2 alone, **(b)** NSP5 fused to a V5 tag alone, and **(c)** their co-expression in RVD, RVF and RVG. **d)** Merged immunofluorescence images of MA/cytBirA cells co-expressing Flag-VP2 with NSP5 (first, third, and fifth columns) or NSP5ΔT (second, fourth, and sixth columns) fused to a BAP tag. Immunofluorescence images of LMH cells expressing a tail deletion of NSP5 fused to a V5 tag of RV species D, F, and G alone **(e)** or in co-expression with its respective Flag-VP2 **(f)**. At 16 hpt, the cells were fixed and immunostained to detect NSP5 or NSP5ΔT (StAv, green or anti-V5, green) and VP2 (anti-Flag, red or cyan). Nuclei were stained with DAPI (blue). The scale bar is 10 µm. The white, red, and yellow arrows point to globular VLS, nuclear inclusions, and aggregated proteins, respectively. The discontinuous yellow lines mark the nucleus as determined by DAPI staining.

The predicted ordered region of NSP5, termed the tail, is found at the N-terminus of RVD and RVF and at the C-terminus of RVA, RVB, RVC, RVG, RVH, RVI, and RVJ, respectively. It has been shown to play a predominant role in VLS formation with NSP2 (65). Moreover, the deletion of the NSP5 tail (NSP5ΔT) impairs VLS formation induced by VP2 in RVA (44). We wondered if the co-expression of NSP5ΔT with VP2 would also impede VLS formation in non-RVA species. In this context (**Fig. 3d**), we compared VLS formation in MA/cytBirA cells by co-expressing Flag-VP2 (cyan) with full-length NSP5 or their respective NSP5 tail deletion fused to the BAP tag (green). Consistent with previous evidence (44), NSP5ΔT-BAP/A impairs VLS formation when co-expressed with Flag-VP2/A. Similar results were obtained for RVC, RVH, RVG, RVI, and RVJ, which also showed impaired VLS formation upon NSP5 tail deletion. Interestingly, co-expression of Flag-VP2 with NSP5ΔT-BAP of RVB leads to irregular filamentous cytosolic structures, contrasting with the diffuse distribution of the two proteins when co-expressed with the full-length NSP5 counterpart. Surprisingly, BAP-ΔTNSP5/D forms VLS with its cognate Flag-VP2. Although BAP-ΔTNSP5/F expressed alone forms nuclear inclusions (**Fig. S3** (65)), co-expression of BAP-ΔTNSP5/F with Flag-VP2/F resulted in the formation of larger and numerous cytosolic VLS as well as nuclear globular inclusions. V5-ΔTNSP5/D and ΔTNSP5/F co-expressed with their cognate Flag-VP2 in LMH cells (**Fig. 3e and f, upper and middle rows**) support the formation of VLS similarly to that in mammalian cells. In contrast (**Fig. 3e and f, bottom row**), the co-expression of NSP5ΔT-V5/G with Flag-VP2/G impairs VLS formation but creates nuclear globular inclusions apparently composed of NSP5ΔT-V5/G.

### The role of the NSP5 tail in its association with VP2 is significant

Since the ordered region of NSP5 plays a critical role in forming VLSs in most RV species studied, we wondered whether this region is also necessary for the direct interaction of NSP5 and VP2. Using pull-down and tripartite GFP assays, we previously demonstrated that NSP5ΔT impairs association with VP2 in RVA (44). In this study, we established a versatile system utilizing bioluminescence resonance energy transfer (BRET) assays between NanoLuc luciferase (NL) fused to VP2 (NL-VP2) and HaloTag fused to NSP5 (HT-NSP5). Essentially, the energy generated from the reaction of the NanoLuc fused protein (NL-VP2) with its substrate (NanoGlo) excites a second fluorescent molecule fused to the HaloTag (HT-NSP5) when in close proximity (**Fig. S4a**), which can be quantified in luminescence units. First, we tested the interaction of NL-VP2 with NSP5-HT of RVA by cell co-expression. As expected (**Fig. 4a**), these two proteins can interact, as indicated by the significant difference (p>0.000001) from experimental controls (HT-NSP5 + NL and HT + NL-VP2). Moreover, the interaction of NL-VP2 with HT-NSP5ΔT compared to HT-NSP5 is significantly impaired. Subsequently, we conducted this assay for RV species B to J by fusing NL to VP2 and HT to NSP5 (**Fig. S4 b-f**) and found that these protein pairs interact significantly with each other (**Fig. 4b-i**). We also assessed the interaction in RV species B to J of NL-VP2 with NSP5 tail deletion fused to HT. We found that this interaction is significantly impaired for all RV species tested, with the exception of RVD.

**Figure 4.**
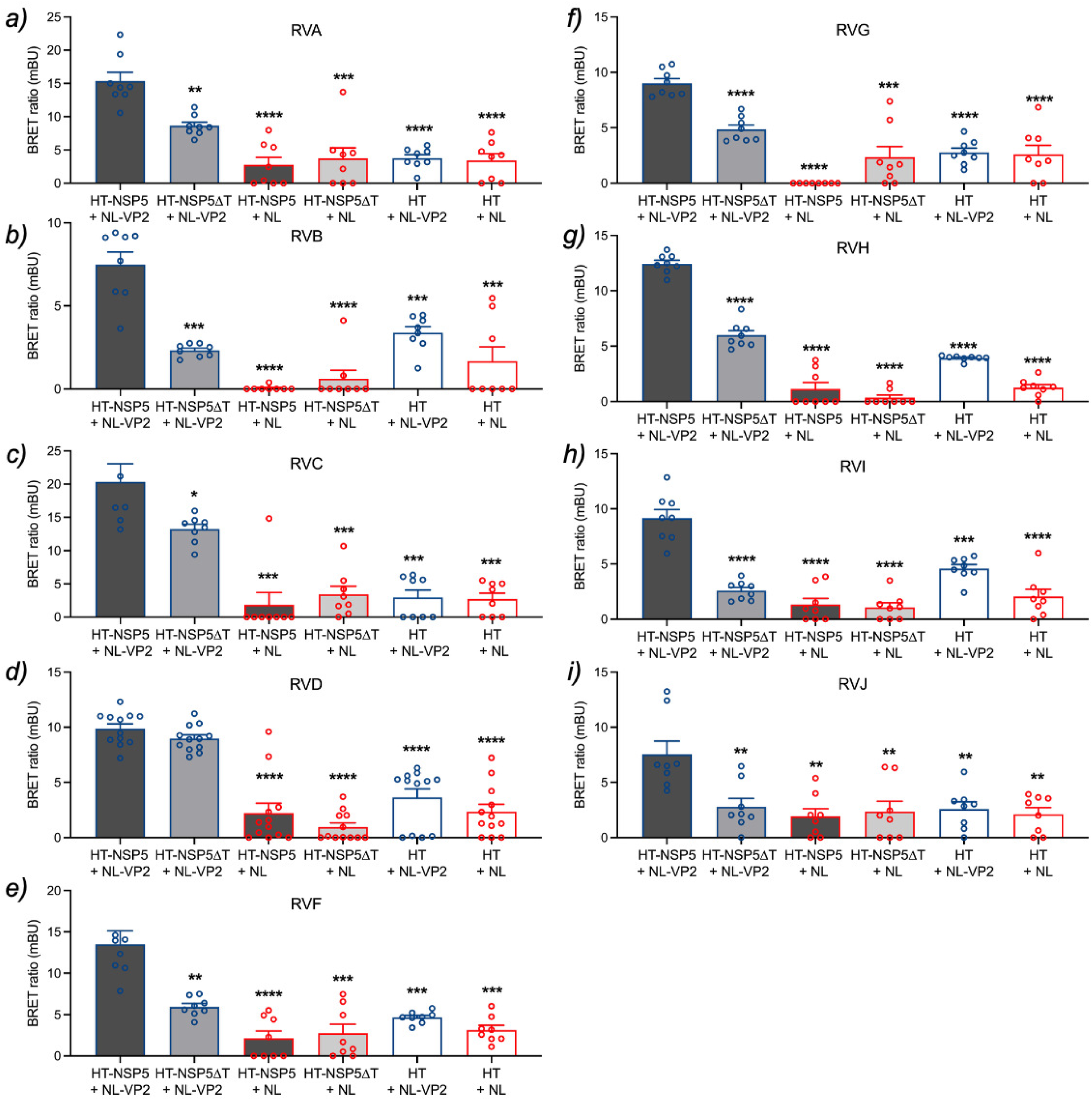
Quantification of VP2 and NSP5 association across RV species A to J. A bioluminescence resonance energy transfer (BRET) assay was conducted to examine the association of NanoLuc luciferase (NL) fused to VP2 with HaloTag (HT) fused to full-length NSP5 or NSP5ΔT for RV species A **(a)**, B **(b)**, C **(c)**, D **(d)**, F **(e)**, G **(f)**, H **(g)**, I **(h),** and J **(i)**. HEK-293T cells were transfected with the indicated pairs for 16 h, then incubated with HaloTag 618 ligand substrate for 6 h followed by the addition of Nano-Glo substrate. Luminescence controls for each protein were included. The result corresponds to the mean ± SEM of three independent experiments. The data were compared to HT-NSP5 with NL-VP2 couple using Brown-Forsythe and Welch ANOVA tests, (*), p<0.05; (**), p<0.01;(***), p<0.001; and (****), p<0.0001.

### VP2 phylogenetic analysis and structural localization of conserved critical residues for VLS formation

Next, we analyzed the evolutionary proximity of VP2 across RV species A to J (**Fig. 5a**) by comparing their CDS available in the database to form pairs of RV species having common ancestors. Similarly, as described for NSP5 and NSP2(65), VP2 presented closer topologies between RV species categorized into two main groups composed of RV species A, C, D, and F, and RV species B, G, H, I, and J. In this context, RVA is closer to RVC, RVF is closer to RVD, RVB is closer to RVG, and RVH is closer to RVJ, where RVI is the most distant but closely related to RVH and RVJ.

**Figure 5.**
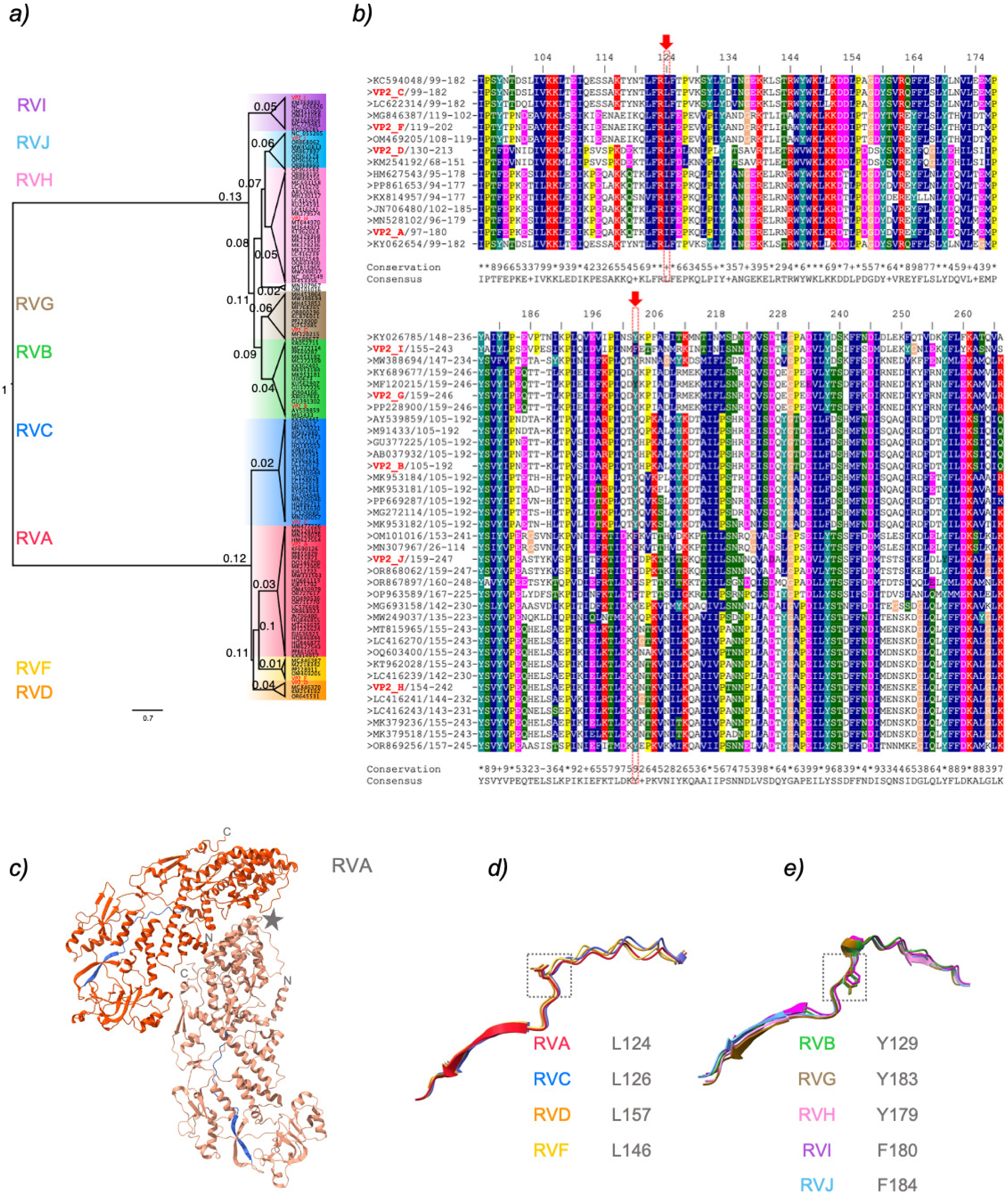
Sequence likelihood of VP2 from RV species A to J. **a)** Maximum likelihood tree illustrating phylogenetic relationships among the VP2 of RV species A to J. The red label corresponds to the CDS used in this study, followed by a letter associated with its RV species. Each RV species has a colored panel: RVA, red; RVB, green; RVC, blue; RVD, orange; RVF, yellow; RVG, brown; RVH, pink; RVI, purple; and RVJ, light blue. The scale is 0.7 substitutions per nucleotide. **b)** Multiple sequence alignment of VP2 from RV species A to J derived from the VP2 phylogenetic tree. GenBank accession numbers are indicated. The conserved residues are labeled according to the ClustalX classification, where blue is hydrophobic, red is positively charged, pink is negatively charged, green is polar, light orange is glycine, yellow is proline, cyan is aromatic, and white is unconserved. The first and last residues of each RV protein region are indicated. The red arrows point to conserved leucine (L) in RV species A, C, D, and F or aromatic residues (F or Y) in RV species B, G, H, I, and J, respectively. **c)** AlphaFold3 prediction of dimeric VP2 of RVA (monomer type A, dark orange, and monomer type B, light orange). The region from 97 to 180 is highlighted in blue for both monomers. A grey star indicates the five-fold axes. N- and C-termini for each monomer are marked. **d)** Overlap of VP2 region 97-180 of RVA with RV species C (blue), D (orange), and F (yellow) in the left panel. A grey dashed open box indicates conserved leucine. Overlapping VP2 regions of RV species B (green), G (brown), H (pink), I (purple), and J (light blue) are shown in the right panel, corresponding to the equivalent VP2 region 97-180 of RVA. Conserved aromatic residues Y or F are highlighted by a grey dashed open box. The values of the predicted local distance difference test (pLDDT) are >70 for all the model predictions.

We previously described (44) that a highly conserved L124 present in VP2 in RVA strain SA11 is essential for viroplasm and VLS formation and its association with NSP5 and RV replication (4). An equivalent motif for L124 is also found in RVC, RVD, and RVF (**Fig. 5b, top panel**), corresponding to VP2 L126, L157, and L146, respectively. However, no equivalent primary sequence with conserved leucine was detected for RVB, RVG, RVH, RVI and RVJ (data not shown). Next, we localized the RVA VP2 region for L124 in the previously described tertiary structure for RVA VP2 strain RRV (69), corresponding to amino acid residues 94 to 180 (**Fig. 5c, blue region**), where L124 is found in a loop preceding a beta-sheet. Using AlphaFold3, we superimposed the VP2 RVA tertiary structure with the predicted VP2 structures for RV species B to J. Consistent with the above alignment, L126 of RVC, L157 of RVD, and L146 of RVF completely overlapped with the L124 of RVA (**Fig. 5d**). Moreover, we found that the predicted structures of RVB, RVG, RVH, RVI, and RVJ overlap with region 97 to 180 of VP2 RVA (**Fig 5e**), having a loop preceding a beta-sheet. Interestingly, the sequence alignment of the loop in these RV species showed conserved aromatic residues corresponding to tyrosine or phenylalanine (**Fig. 5b**, **bottom panel**). Thus, the Y129 of RVB, Y183 of RVG, Y179 of RVH, F180 of RVI, and F184 of RVJ overlap with the respective predicted loop at an equivalent position as L124 in VP2 RVA.

### Conserved VP2 residue is essential for VLS formation

We hypothesized that a conserved residue in VP2 of RV species B to J is necessary for VLS formation, similar to what has been described for L124 in RVA. Accordingly, a point mutation substituting L124 with a non-bulky amino acid like alanine (L124A) resulted in impaired VLS formation (44). To investigate this, we expressed Flag-VP2 constructs with alanine substitutions at conserved residues across RV species A to J. These mutant proteins migrated at their predicted molecular weights when expressed in MA104 cells (**Fig. 6a**). Notably, Flag-VP2(Y129A)/B showed weak expression, although it migrated at the correct molecular weight.

**Figure 6.**
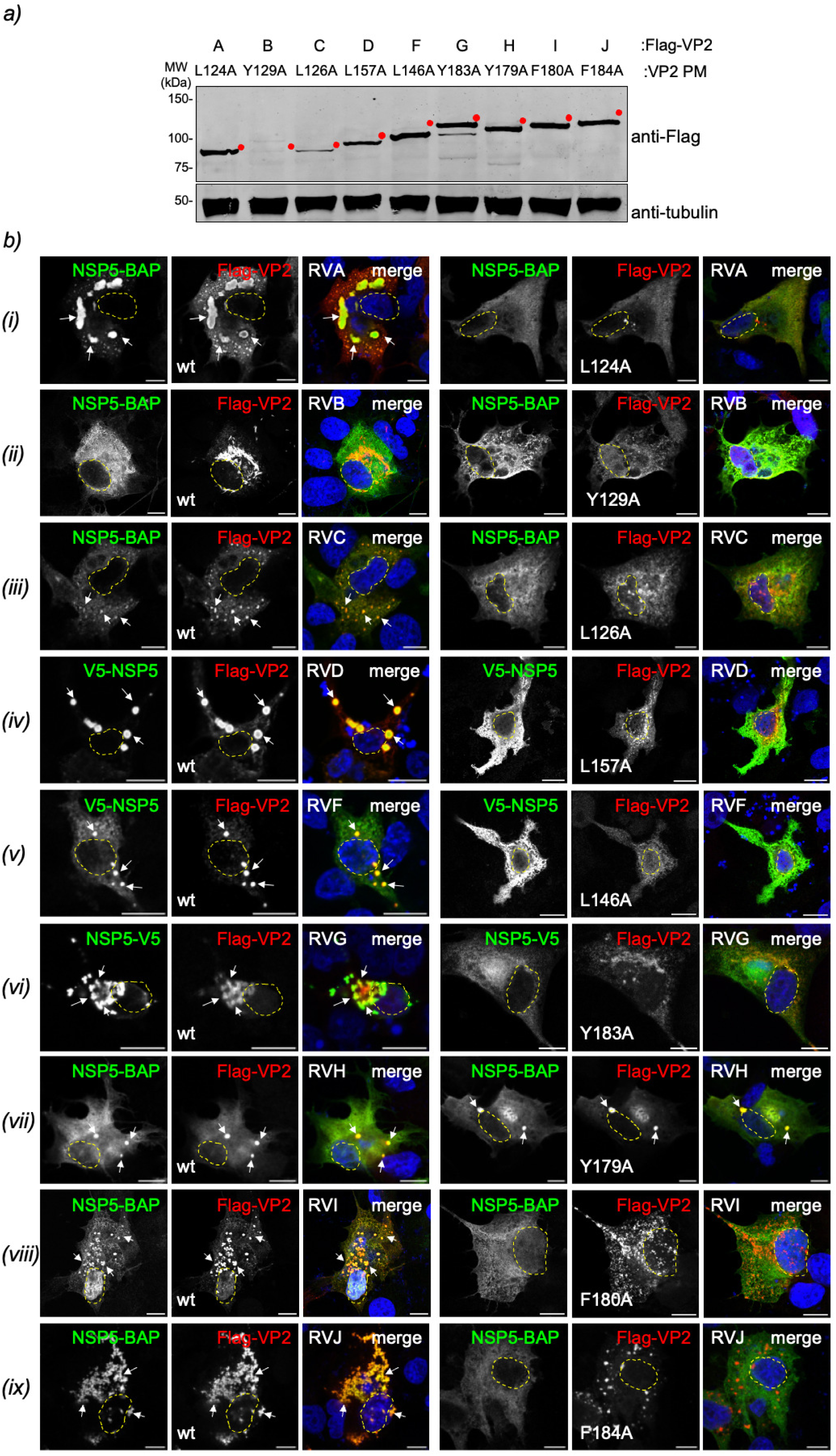
Point mutations to conserved residues in VP2 of RV species A to J impair VLS formation. **a)** Immunoblotting of cellular lysates from MA104 cells expressing Flag-VP2 of RV species A to J with the indicated point mutation. The membrane was incubated with mouse mAb anti-Flag for the detection of Flag-VP2 (top panel) and mouse anti-tubulin (bottom panel) for the detection of alpha-tubulin as a loading control. The red dot indicates the predicted molecular weight. **b)** Immunofluorescence images comparing VLS formation of cells co-expressing NSP5 with wtVP2 (left panel) or VP2 containing the indicated point mutation (right panel) of RV species A to J. At 16 hpt, the cells were fixed and co-stained to detect NSP5-BAP (StAv, green) or V5 tag fusion to NSP5 (anti-V5, green, RV species D, F, and G) with Flag-VP2 (anti-Flag, red). The third and sixth columns correspond to merged images. The scale bar is 10 µm. VLS formation was detected in MA/cytBirA cells for RV species A (*i*), B (*ii*), C (*iii*), H (*vii*), I (*viii*), and J (*ix*), while LMH cells were utilized for detection in RV species D (*iv*), F (*v*), and G (*vi*), respectively. The white arrows point to globular VLS. Nuclei are outlined with a dashed yellow line as determined by DAPI staining.

We then assessed VLS formation by immunofluorescence microscopy in MA/cytBirA cells co-expressing NSP5-BAP with either wild-type (wt) Flag-VP2 or Flag-VP2 point mutants from RVA, RVB, RVC, RVH, RVI, and RVJ (**Fig. 6b, rows *i*, *ii*, *iii*, *vii*, *viii*, and *ix***). Consistent with previous findings, the co-expression of NSP5-BAP with RVA Flag-VP2 (L124A) failed to support VLS formation (**Fig. 6b, row *i***). Similarly, VLS formation was impaired when co-expressing Flag-VP2 point mutants of RVC, RVI, and RVJ with their respective NSP5-BAP, in marked contrast to the VLS observed with their wild-type (wt) counterparts. In RVB, the co-expression with wt Flag-VP2 with its cognate NSP5-BAP did not lead to VLS formation; therefore, it was expected that the Y129A mutant would fail to form VLS (**Fig. 6b row *ii***). In contrast, Flag-VP2 (Y179A) co-expression with NSP5-BAP of RVH supported VLS formation (**Fig. 6b, row *vii***).

The Flag-VP2 point mutants from avian RVD, RVF, and RVG were tested in LMH cells to provide a suitable host environment (**Fig. 6 b, rows *iv*, *v*, and *vi***). While the co-expression of wtFlag-VP2 with its cognate NSP5 fused to V5 supported VLS formation in these RV species, the respective alanine mutants – L157A of (RVD), L146A (RVF), and Y183A (RVG) – failed to form VLSs.

### VLS morphology is modulated by NSP5, NSP2, and VP2

We previously reported that the co-expression of cognate NSP5 with NSP2 leads to the formation of globular VLSs for RVA, RVB, RVD, RVG, and RVI. In contrast, RVC forms filamentous VLS, while RVH and RVJ do not form VLS (**Table 2**) (65). In this study, we observed that co-expression of NSP5 with VP2 resulted in the formation of globular VLSs in all RV species except for RVB. We next investigated whether NSP2 expression could influence the morphology of VP2-induced VLSs (**Table 2**). For this purpose (**Fig. 7a**), we compared the morphology of VLS formed by the co-expression of NSP5 and VP2 [VLS (VP2)i] with those formed by the co-expression of NSP5, NSP2, and VP2 [VLS (VP2+NSP2)i]. We found that under the latter condition, VLSs exhibit a globular morphology across all RV species except for RVC, which retained a filamentous morphology. Notably, globular VLS (VP2+NSP2)i facilitated the recruitment of NSP2 in RVH and RVJ and the recruitment of VP2 in RVB.

**Figure 7.**
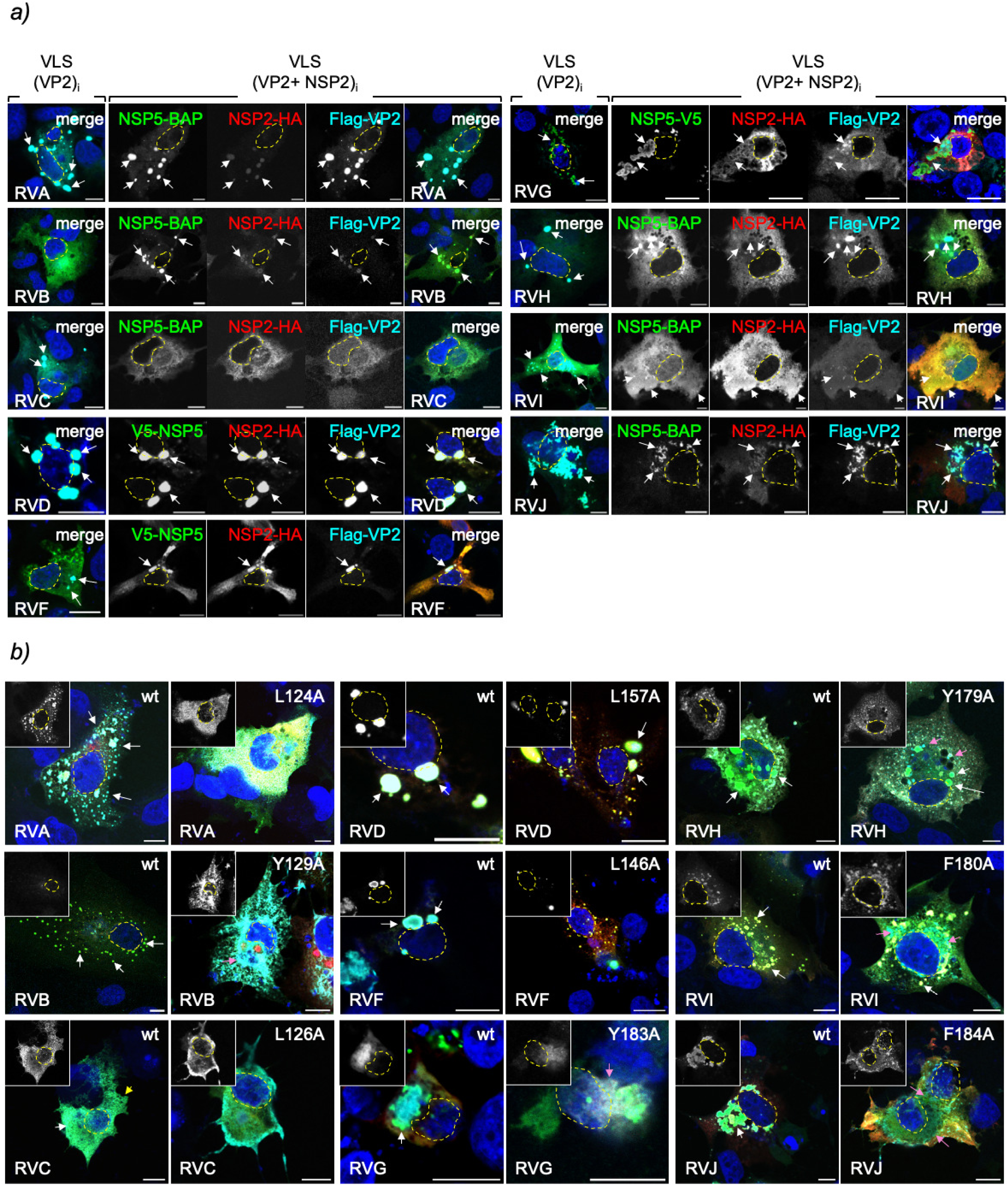
Morphology of VLS composed of NSP5 with VP2 and NSP2 across RV species A to J. **a)** Immunofluorescence images of cells co-expressing BAP-tagged or V5-tagged NSP5 with Flag-VP2 in the absence (VLS(VP2)i) and presence (VLS(VP2+NSP2)i) of NSP2-HA for RV species A to J. At 16 hpt, the cells were fixed and stained for detection of BAP-tagged NSP5 (RVA, RVB, RVC, RVH, RVI, and RVJ; StAv, green) or V5-tagged NSP5 (RVD, RVF, and RVG; anti-V5, green), NSP2-HA (anti-HA, red) and Flag-VP2 (anti-Flag, cyan). Nuclei were stained with DAPI (blue). The scale bar is 10 µm. The white arrows point to globular VLSs. The nuclei were labeled with a yellow dashed line as determined by DAPI staining. **b)** Immunofluorescence images comparing VLS composed of BAP-tagged or V5-tagged NSP5, NSP2-HA, and Flag-VP2 wt or its corresponding point mutation. The cells were fixed at 16 hpt and stained for the detection of BAP-tagged NSP5 (RVA, RVB, RVC, RVH, RVI, and RVJ; StAv, green) or V5-NSP5 (RVD, RVF, and RVG; anti-V5, green), NSP2-HA (anti-HA, red), and Flag-VP2 (anti-Flag, cyan). Nuclei were stained with DAPI (blue). The scale bar is 10 µm. The image corresponds to the merged image. The top left corner of each image corresponds to Flag-VP2 immunostaining. The white, yellow, and pink arrows point to globular VLS, filamentous VLS, and dispersed VP2, respectively. The nuclei were labeled with a yellow dashed line as determined by DAPI staining. In both experiments, VLS formation was detected in MA/cytBirA cells for RVA, RVB, RVC, RVH, RVI, and RVJ, while LMH cells were used for RVD, RVF, and RVG.

**Table 2.** Summary of VLS formation through the co-expression of NSP5 with either NSP2, VP2, or both.

| RV species | VLS(VP2)<sub>i</sub> |  |  |  | VLS(NSP2)<sub>i</sub> formation<sup>c</sup> |  | Morphology VLS (VP2+NSP2)<sub>i</sub> |
| --- | --- | --- | --- | --- | --- | --- | --- |
|  | VLS formation<sup>a</sup> |  | NSP5-VP2 interaction<sup>b</sup> |  |  |  |  |
|  | NSP5 | NSP5ΔT | NSP5 | NSP5ΔT | NSP5 | NSP5ΔT |  |
| A | + | - | + | - | + | - | globular |
|  | (globular) |  |  |  | (globular) |  |  |
| B | - | - | + | - | + | - | globular |
|  |  |  |  |  | (globular) |  |  |
| C | + | - | + | - | + | - | filamentous |
|  | (globular) |  |  |  | (filamentous) |  |  |
| D | + | + | + | + | + | + | globular |
|  | (globular)<sup>d</sup> | (globular) |  |  | (globular) | (globular) |  |
| F | + | + | + | - | + | + | globular |
|  | (globular) | (globular)<sup>e</sup> |  |  | (globular) | (globular)<sup>e</sup> |  |
| G | + | - | + | - | + | - | globular |
|  | (globular) |  |  |  | (globular)<sup>c</sup> |  |  |
| H | + | - | + | - | - | - | globular |
|  | (globular) |  |  |  |  |  |  |
| I | + | - | + | - | + | - | globular |
|  | (globular) |  |  |  | (globular) |  |  |
| J | + | - | + | - | - | - | globular |
|  | (globular) |  |  |  |  |  |  |

*a)* Confirmed by co-immunostaining for the detection of both VLS components, NSP5 and
VP2 (transfection ratio 1:2= NSP5:VP2).
*b)* Determined by BRET assay.
*c)* Described by Lee et al., 2024 (65)
*d)* Forms VLS only in chicken epithelial LMH cells.
*e)* Forms nuclear inclusions
*f)* +, positive; -, negative.

Additionally, we analyzed the impact of VP2 point mutants on VLS(VP2+NSP2)i morphology (**Fig. 7b**). As previously described by Buttafuoco et al. (44), the Flag-VP2(L124A) disrupted VLS(VP2+NSP2)i in RVA. The corresponding VP2 point mutants in other RV species also disrupted VLS formation. This was evidenced by the appearance of small, punctuated VLSs lacking VP2 in RVD and RVF and by irregular VLSs with cytosolic dispersion of VP2 in RVB, RVG, RVH, and RVI. Interestingly, VLS formation was impaired entirely in RVC and RVJ, resulting in the loss of their characteristic filamentous and globular morphologies, respectively.

### Heterologous formation of VP2-induced VLS among RV species

We previously described that NSP5 and NSP2 from closely related RV species couples can be interchanged, forming heterologous VLSs (65). This was the case for RVA with RVC, RVB with RVG, and RVD with RVF. Since VP2 has the same phylogenetic ancestor distribution as NSP5 and NSP2 (**Fig. 5a**), we wondered if heterologous VLS could be formed between closely related couples of NSP5 and VP2. In this context (**Fig. 8a, top panel**), we co-expressed NSP5-BAP with Flag-VP2 of RVA and RVC in all four interspecies combinations of NSP5 and VP2 (A/A, C/C, A/C, and C/A). We found that all the combinations support VLS formation, although they have diverse sizes, from large (NSP5/RVA with VP2/RVA) to punctuated (NSP5/RVC with VP2/RVA). The combinations of NSP5 and VP2 of RVB and RVG lacked the ability to form heterologous VLSs in both mammalian (**Fig. 8a, middle panel**) and avian (**Fig. 8b, top panel**) cells. Interestingly, RVH with RVJ (**Fig. 8a, bottom panel**) and RVD with RVF (**Fig 8b, bottom panel**) interchanged their respective NSP5 and VP2, forming heterologous VLS in all four combinations.

**Figure 8.**
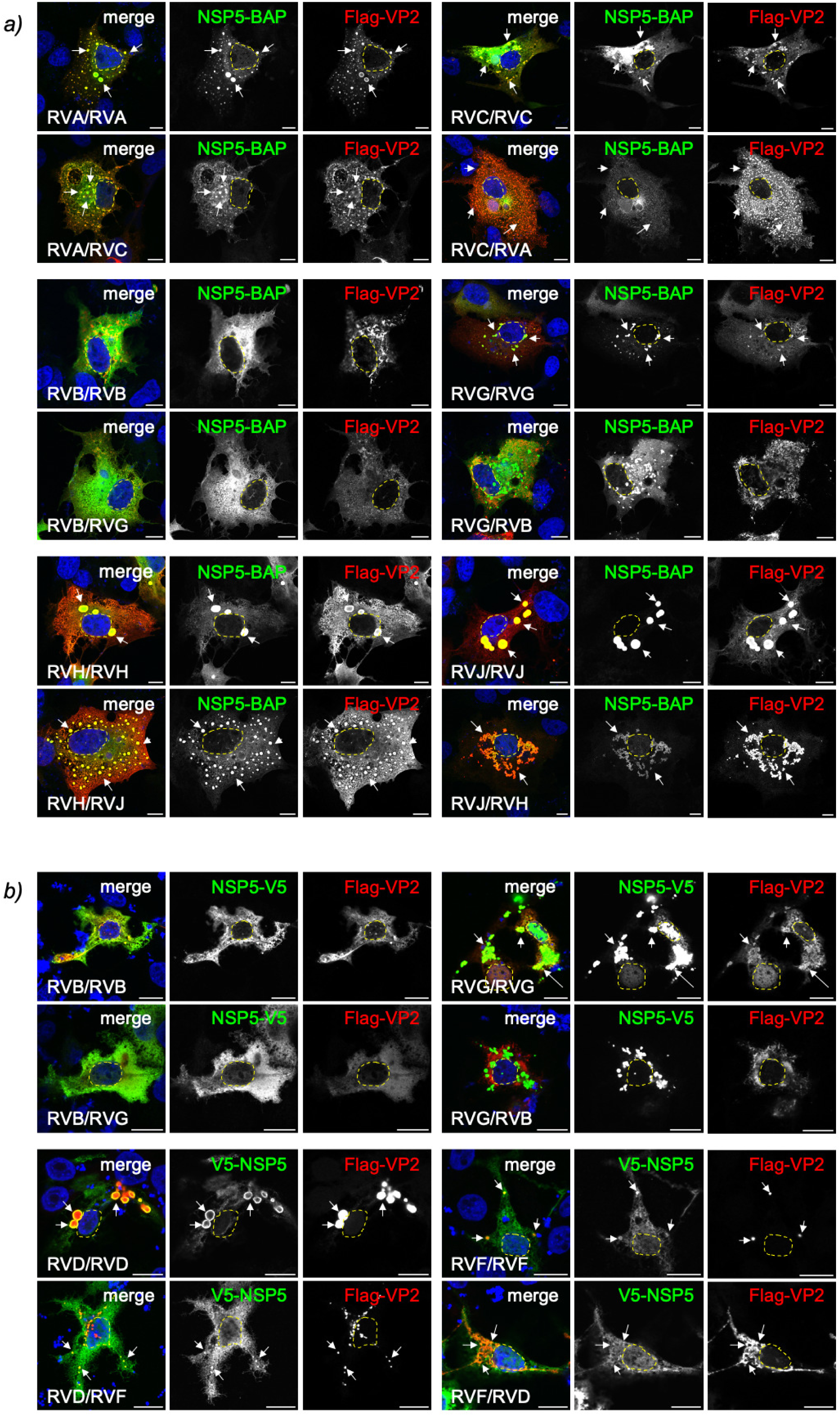
Heterologous formation of VLS by NSP5 and VP2 from closely related RV species. **a)** Immunofluorescence images of MA/cytBirA cells co-expressing BAP-NSP5 and Flag-VP2 from closely related RVA and RVC (top panel), RVB and RVG (middle panel), and RVH and RVJ (bottom panel) in all indicated combinations. After fixation, the cells were immunostained for the detection of NSP5 (StAv, green) and VP2 (anti-Flag, red). The nuclei were stained with DAPI (blue). **b)** Immunofluorescence images of chicken LMH cells co-expressing V5-tagged NSP5 with Flag-VP2 from closely related RVB and RVG (top panel), and RVF and RVD (bottom panel). After fixation, the cells were immunostained for the detection of NSP5 (anti-V5, green) and VP2 (anti-Flag, red). The nuclei were stained with DAPI (blue). In all images, the indication at the bottom left corner corresponds to the RV species of NSP5 and VP2, respectively. The scale bar is 10 µm. The white arrows point to globular VLS. The yellow lines label the nucleus position.

### Chimeric VP2 RVB harboring the N-terminal region of VP2 RVG forms VLS

Previously, we demonstrated that the co-expression of NSP5 and NSP2 of RVB formed globular VLS, with NSP5 also capable of forming heterologous VLS with NSP2 of RVG (65). However, co-expression of NSP5 of RVB did not lead to VLS formation with either homologous VP2 of RVB or heterologous VP2 of RVG, although VP2 of RVG supported VLS formation with its cognate NSP5 in avian cells. We next reasoned that VP2 of RVB might differ from VP2 of other RV species because it lacks an unstructured N-terminal region (**Fig. 1c**). We therefore hypothesized that the unstructured N-terminal region of VP2 plays a functional role in forming globular VLS. To investigate this, we used AlphaFold3 to compare the predicted tertiary structure of VP2 of RVB with that of its close relative, VP2 of RVG. The first common structural element identified was a beta-sheet starting at valine 85 in VP2 of RVB and aspartic acid 138 in VP2 of RVG. We then replaced the N-terminal region of VP2 of RVB *in silico* with amino acids 1 to 137 from VP2 of RVG (**Fig. 9a**), generating a chimeric VP2/G-B. This chimera was predicted to contain a disordered N-terminal region similar to that of VP2/G while preserving the apical, central, and dimerization regions of VP2 of RVB. We then constructed and expressed the chimeric VP2/G-B fused to a Flag-tag, which migrated at the predicted molecular weight (**Fig. 9 c and d**).

**Figure 9.**
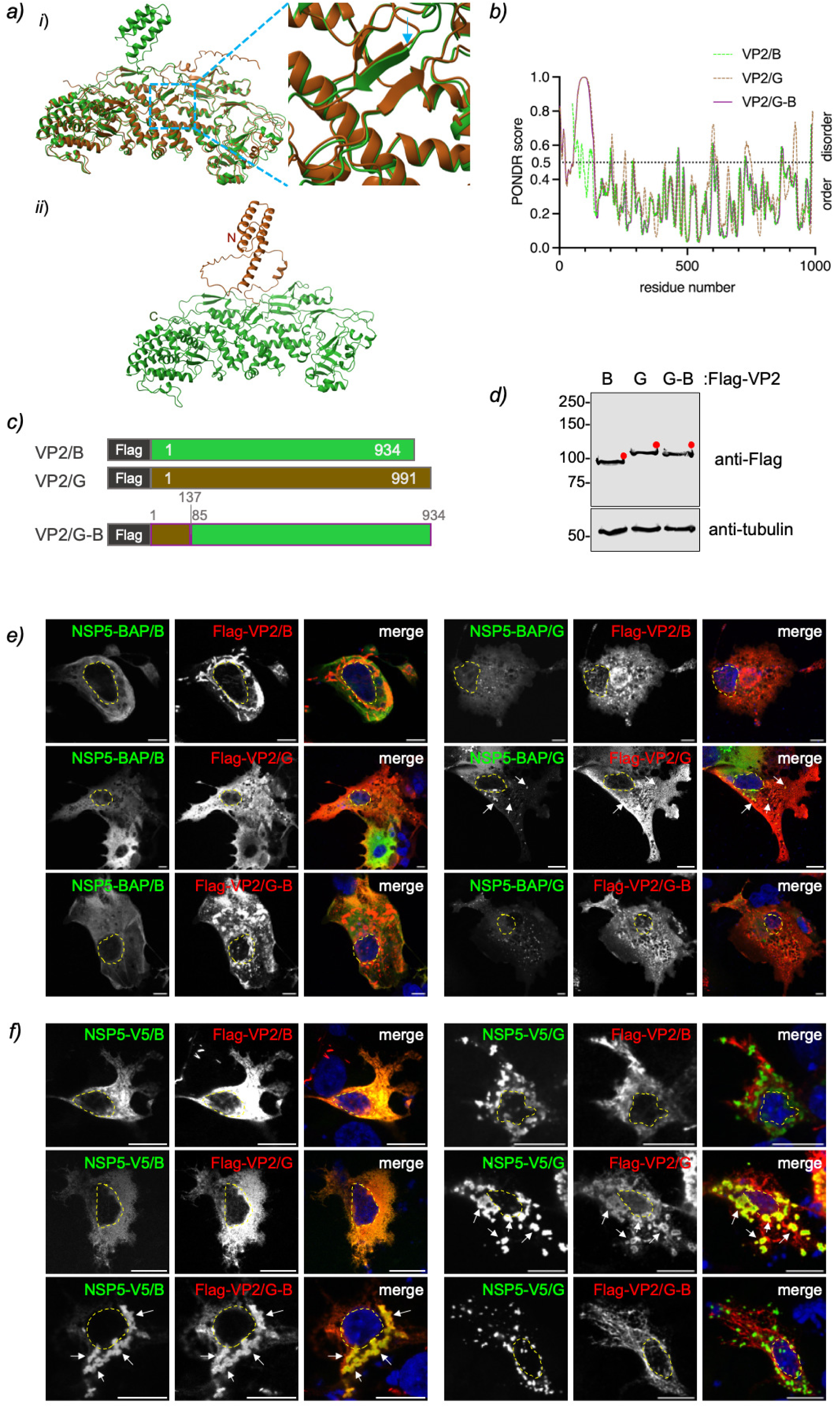
The disordered N-terminal region of VP2 is crucial to VLS formation. **a)** AlphaFold3 prediction for the design of chimeric VP2/G-B. (*i*) Overlap of predicted monomeric structures of VP2/B (amino acids 1 to 934, green) and VP2/G (amino acids 101 to 991, brown). The open dashed blue frame corresponds to the magnified image on the right, which shows the initial overlapping structures between VP2/B and VP2/G. The blue arrow points to V85 in VP2/B and D138 in VP2/G. (*ii*) The predicted monomeric structure of chimeric VP2/G-B is displayed in the VP2/G N-terminal region (amino acid residues 1 to 137, brown) alongside VP2/B from residue 85 to 934 (green). **b)** Plot comparing IDR predictions of VP2 species B and G and chimeric VP2/G-B. **c)** Schematic representation of Flag-VP2/B and G, along with chimeric Flag-VP2/G-B constructs. The chimeric Flag-VP2/G-B contains the N-terminal region 1 to 137 of VP2/G and the region 85 to 934 of VP2/B. **d)** Immunoblotting of cellular lysates from MA104 cells expressing Flag-VP2/B, Flag-VP2/G, and chimeric Flag-VP2/G-B. The membrane was incubated with anti-Flag (top panel) and anti-tubulin (bottom panel) as a loading control. The red dots indicate the predicted molecular weight of each protein. Immunofluorescence images of co-expression of NSP5/RVB (left panel) or NSP5/RVG (right panel) with Flag-VP2/B (first row), Flag-VP2/G (second row), or chimeric Flag-VP2/G-B (third row). At 16 hpt, MA/cytBirA **(d)** or LMH **(e)** cells were fixed and stained for detection of NSP5 (StAv **(d)**, anti-V5 **(e)**, green) and VP2 (anti-Flag, red). The scale bar is 10 µm. The white arrows point to globular VLS. The nuclei are outlined with a dashed yellow line as determined by DAPI staining (blue).

Next, we co-expressed BAP-tagged or V5-tagged NSP5/B (**left panel**) or NSP5/G (**right panel**) with Flag-tagged VP2/B, VP2/G, or the chimeric VP2/G-B in mammalian (**Fig. 9e**) or avian (**Fig. 9f**) cells. As expected, VLS formation was observed for NSP5 and VP2 of RVG in both cell types. In mammalian cells, no other protein combinations supported VLS formation. Interestingly, in avian cells, co-expression of NSP5/B with the chimeric VP2/G-B supported VLS formation, but not when NSP5/G was co-expressed with VP2/G-B.

## Discussion

Understanding the RV life cycle, especially the assembly of the virions, is essential for addressing its spread. RV comprises nine species, from A to J, that infect a broad range of mammals and birds. Of note, two new RV species (RVK and RVL) were recently added by the ICTV, although they were not included in this study. RVA viroplasms are cytosolic globular inclusions that support virus genome replication, sorting, and packaging in newly assembled viral cores. The lack of research tools makes studying the life cycle of non-RVA species challenging. We recently tackled this significant issue by extrapolating the role of ortholog proteins responsible for VLS formation to non-RVA(65). In this way, we described the ability of NSP5 to form VLS in co-expression with NSP2 in certain RV species, such as RVB, RVD, RVF, RVG, and RVI. Similarly, in the current study, we investigated VLS formation across RV species A to J by co-expressing NSP5 with VP2 using confocal immunofluorescence microscopy. We found that VLS can form under these conditions in RVA, RVC, RVD, RVF, RVG, RVH, RVI, and RVJ, but not in RVB. These results contrast with those obtained from our previous study, particularly for RVB, RVH, and RVJ. In this context, it appears that for certain RV species, like RVB, NSP2 is essential for VLS formation. In contrast, for other RV species, such as RVH and RVJ, where NSP2 is dispensable, VP2 is essential. Like for RVA, NSP2 and VP2 had a complementary role in the VLS formation of RVC, RVD, RVF, RVG, and RVI. We also show that NSP5 and VP2 directly interact, as denoted by the BRET assay, indicating a correlation of their association with VLS formation. These experiments therefore confirm the significant role of VP2 in viroplasm formation.

Similar to how VLSs form with NSP2 (16, 61, 65), we also demonstrate that the tail region of NSP5 is essential for both its interaction with VP2 and the induction of VLSs across multiple RV species, including RVA, RVC, RVG, RVH, RVI, and RVJ (**Table 2**). NSP5ΔT/B exhibited an impaired association with VP2 compared to NSP5/B, although neither formed VLS with VP2. However, it is important to note that NSP5/B forms VLS with NSP2, while NSP5ΔT/B does not. We previously described that the deletion of the ordered region of NSP5 in RVD and RVF, located at their N-terminus instead of the C-terminus as in other studied RV species, does not affect VLS formation induced by NSP2 (65). Similarly, the co-expression of ΔTNSP5 with VP2 of RVD or RVF enhances VLS formation in mammalian cells while consistently forming VLS in chicken epithelial cells. Notably, the VLS of ΔTNSP5 with VP2 of RVF also formed nuclear globular inclusions in both cell types, seemingly composed solely of NSP5. Therefore, the nuclear translocation of ΔTNSP5/F influences its cytosolic interaction with VP2, which is consistent with the decreased binding of these two proteins in the BRET assay.

The viroplasms are complex structures composed of several viral proteins, each potentially contributing to viroplasm morphology. We previously demonstrated that RVA VLS induced by either NSP2 or VP2 can recruit other viral proteins (25, 32, 44, 66). Here, we determined that VLS induced by VP2 could incorporate NSP2 for RV species A to J. Interestingly, when NSP2, NSP5, and VP2 from various RV species were expressed, most formed VLSs with globular shape. However, RVC was an exception, producing filamentous VLSs, similar to those induced by RVC NSP5 and NSP2. This result suggests that NSP2 plays a major role, over other RVC proteins, in determining the morphology of RVC VLSs. Likewise, VLS composed of NSP5, NSP2, and VP2 of RVB presents a globular morphology similar to VLS formed by NSP5/B and NSP2/B, indicating that VP2 is recruited to VLS by at least interacting with NSP5. In contrast, VLS induced by VP2 from RVH and RVJ permitted the recruitment of NSP2, maintaining their globular morphology (**Table 2**). We previously demonstrated that the association of the respective NSP5 and NSP2 of RVH and RVJ is weak or not detectable (65). Here, we show that NSP5 and VP2 from these RV species interact and form VLSs, suggesting that VLSs composed of NSP5, NSP2, and VP2 arise either through direct interaction of both NSP5 and NSP2 with VP2, or that VP2 enhances the otherwise weak association between NSP5 and NSP2.

Our earlier findings showed that a conserved residue in VP2 RVA, L124, is necessary for the formation of VLS as well as for maintaining globular morphology and the ability of viroplasms to replicate (44). Here, we found that this conserved residue occupies a similar tertiary position in RV species B through J, as a leucine for RVC, RVD, and RVF, and as an aromatic residue, tyrosine, for RVB, RVG, and RVH, and phenylalanine for RVI and RVJ. Indeed, substituting this conserved residue with alanine disrupts VLS formation in RV species B to J, whether the VLS are induced by VP2 or formed by a combination of NSP5, NSP2, and VP2. The resulting disrupted VLSs showed two distinct patterns: in RVA, RVC, and RVJ, the proteins were completely dispersed throughout the cytosol, while RVB, RVD, RVF, RVG, RVH, and RVI, the VLSs were smaller and mainly composed of NSP5 and NSP2, with VP2 dispersed in the cytosol. These observations suggest that this conserved residue plays a critical structural role in VLS formation across RV species A to J.

Heterologous VLS formation is also observed with NSP5 and VP2 from closely related RV species, such as RVA with RVC, RVF with RVD, and RVH with RVJ, suggesting that genetic reassortment among these RV species may be possible in principle. In this sense, the NSP5 and NSP2 of closely related RV species, RVA with RVC and RVD with RVF, can also be interchanged (65). Since RVH and RVJ do not form VLS with NSP5 and NSP2, it remains unclear whether they can be interchangeable. However, we now demonstrate that NSP5 and VP2 of RVH and RVJ can form interspecies VLS. The formation of triple VLS involving NSP5, NSP2, and VP2 among RVH and RVJ suggests reassortment may also occur. An interesting case involves RVB and RVG, which previously showed the ability to form heterologous VLS between NSP5 and NSP2. However, this was not the case for NSP5 and VP2. In this context, we demonstrate that the absence of a disordered N-terminal region on VP2/B hinders VLS formation, whereas replacing its N-terminal with that of VP2/G permits VLS formation with NSP5/B but not with NSP5/G. This finding indicates that reassortment in these RV species will depend on the interchange of heterologous NSP5 and NSP2. Nonetheless, we cannot rule out the possibility that a natural recombination of VP2 of RVB with its closely related VP2 RVG could lead to the acquisition of a disordered N-terminal region. In this study, we also describe for the first time that the disordered region of VP2 not only plays a role in the association with replication intermediates VP1 and VP3 in the core virion (58, 60, 62, 70) but also in the formation of VLS and, by extension, probably of viroplasms.

RV reverse genetics has been established only for certain RVA strains, such as simian SA11 (71), porcine OSU (72), and human KU (73) and is not available for other RVA strains and non-RVA species. Understanding viroplasms is crucial for applying reverse genetics to non-RVA species, as the co-expression of proteins like NSP5 and NSP2 significantly enhances the recovery of recombinant rotaviruses (74). For future experiments exploring reverse genetics in other RV species, it appears that for RVH and RVJ, the co-expression of NSP5 and VP2 will be favored, instead of NSP5 and NSP2, for the successful recovery of recombinant virus.

## Materials and Methods

### Cells and viruses

MA104 (embryonic rhesus monkey kidney, ATCCCRL-2378, RRID: CVCL_3845) cells were cultured in Dulbecco’s modified Eagle’s medium (DMEM, Gibco BRL) supplemented with 10% fetal calf serum (FCS, AMIMED, Bioconcept, Switzerland) and penicillin (100 U/ml)-streptomycin (10 µg/ml). MA/cytBirA (25) were cultured in DMEM supplemented with 10% FCS, penicillin (100 U/ml)-streptomycin (10 µg/ml), and 5 µg/ml puromycin (InvivoGen, France). LMH cells (chicken hepatocellular carcinoma epithelial, ATCCCRL2117) were cultured in Waymouth’s MB572/1 (Sartorius) medium supplemented with 10% FCS and penicillin (100 U/ml)-streptomycin (100 µg/ml). HEK-293T (human embryonic kidney, ATCCCRL-3216) cells were cultured in DMEM supplemented with 10% FCS and penicillin (100 U/ml)-streptomycin (10 µg/ml).

The recombinant vaccinia virus encoding T_7_ RNA polymerase (strain vvT7.3) was amplified as previously described (75).

### Antibodies and reagents

Guinea pig anti-VP2 was described previously (66). Mouse monoclonal (mAb) anti-tubulin (clone B5-1-12) and mouse mAb anti-Flag (clone M2) were purchased from Merck. AlexaFluor594 anti-HA.11 (clone 16B12) and AlexaFluor 647 anti-Flag Tag (clone L5) were purchased from BioLegend. Mouse mAb-V5 Tag-Dylight 488 was purchased from Invitrogen. Streptavidin-Dylight488 and mouse secondary antibodies conjugated to AlexaFluor 488 or AlexaFluor 594 were purchased from Thermo Fisher Scientific. The secondary antibodies for immunoblot conjugated to IRDye680CW and IRDye800CW were purchased from LI-COR. Mouse mAb anti-NanoLuc^®^ and HaloTag^®^TMRDirect^™^Ligand (Cat# G2991) were purchased from Promega.

### Rotavirus sequences

The sequences of rotavirus NSP5 and NSP2 open reading frames from species B to J used in this study were previously published by Lee et al., 2024 (65). The sequences of the rotavirus VP2 open reading frames from RV species A to J are provided in the supplemental material and **Table 1**.

### Plasmid constructs

The plasmids pCI-NSP5-BAP/A, B, C, D, F, G, H, I and J; pCI-BAP-NSP5/D and F, pCI-NSP5-V5/G, pCI-NSP5ΔT-BAP/A, B, C, G, H, I and J; pCI-BAP-ΔTNSP5/ and F; and pCI-NSP2-HA/A, B, C, D, F, G, H, I and J were described previously (65). pCI-V5-NSP5/D, pCI-V5-NSP5(15–195)/D, pCI-V5-NSP5/F and pCI-V5-NSP5(19–218)/F were obtained by PCR amplification of pCI-NSP5/D and F(65) using specific primers to insert *Mlu*I/V5 tag and *Not*I sites, followed by ligation into those sites in pCI-Neo (Promega). pCI-Flag-VP2/A was obtained by PCR amplification of pCI-HA-VP2(SA11) (44) using specific primers to insert *Mlu*I/Flag tag and *Not*I sites, followed by ligation in those sites in pCI-Neo. pCI-Flag-VP2/B, C, F, and I were obtained by PCR amplification from baculovirus encoding VP2/B, C, F, and I (gently provided by Dr. Daniel Luque, USW, Australia) using specific primers *Mlu*I/Flag tag and *Not*I sites, followed by ligation in those sites in pCI-Neo. pCI-Flag-VP2/D, G, H, and J were obtained from digestion with *Mlu*I and *Not*I of DNA synthetic segments (GenScript, Netherlands, **Table S2**) encoding for the respective Flag-VP2 and ligation on those sites in pCI-Neo.

The plasmids pCI-HaloTag-NSP5/A, B, C, D, F, G, H, I, and J were obtained digestion of *Mlu*I and *Not*I restriction enzymes of their respective pCI-NSP5(65) and cloned in pCI-HaloTag on those sites. The plasmid pCI-HaloTag was obtained by synthesis of the HaloTag fragment (GenBank: <u>MG867371.1</u>) flanked at 5’- and 3’-ends by *Xho*I and *Mlu*I restriction sites (GenScript, Netherlands) and ligated in those in pCI-Neo (Promega). The plasmids pCI-HaloTag-NSP5(1–178)/A, pCI-HaloTag-NSP5(1–124)/B, pCI-HaloTag-NSP5(1–150)/C, pCI-HaloTag-NSP5(15–195)/D, pCI-HaloTag-NSP5(19–218)/F, pCI-HaloTag-NSP5(1–114)/G, pCI-HaloTag-NSP5(1–151)/H, pCI-HaloTag-NSP5(1–104)/I, and pCI-HaloTag-NSP5(1–136)/J were obtained by digestion with *BspE*I and *Mlu*I restriction enzymes from their respective pCI-NSP5ΔT-BAP (65) and ligated in those sites in pCI-HaloTag-MEB-stop. The plasmid pCI-HaloTag-MEB-stop was obtained by annealing of the oligonucleotides 5’-cgcgtgaattctccggatgagc-3’ and 5’-ggccgctcatccggagaattca-3’, followed by ligation in pCI-HaloTag between *Mlu*I and *Not*I restriction sites. pCI-NanoLuc-Flag-VP2/A, B, C, D, F, G, H, I, and J were obtained from PCR amplification of their respective pCI-Flag-VP2 using a specific primer to insert in frame *Mlu*I and *Not*I restriction sites in Flag-VP2. The PCR fragment was subsequently ligated *Mlu*I/*Not*I sites in pCI-NanoLuc. The plasmid pCI-NanoLuc was obtained by synthesis of the NanoLuc fragment (GenBank: <u>AHH41346.1</u>) flanked at 5’and 3’ ends by *Xho*I and *Not*I restriction sites (GenScript, Netherlands) and ligated in those in pCI-Neo (Promega).

The version of the constructs pCI-Flag-VP2/A, B, C, D, F, G, H, I, and J as well as pCI-NanoLuc-Flag-VP2/A, B, C, D, F, G, H, I, and J harboring VP2 point mutations L124A, Y129A, L126A, L157A, L146A, Y183A, Y179A, F180A, and F184A, respectively, were built by insertion of point mutations using the QuickChange site-directed mutagenesis protocol (Agilent).

The chimeric pCI-Flag-VP2/G-B was obtained by insertion in between *Mlu*I and *Pci*I of pCI-Flag-VP2/B of a synthetic DNA segment (GeneArt Technology, Invitrogen, **Table S2**) containing an in-frame sequence of Flag tag, N-terminal region of VP2/G (region 1-137) and VP2/B region 85 to 163

All the oligonucleotides were obtained from Microsynth AG, Switzerland, and described in **Table S3.**

### AlphaFold predictions

Protein structures of VP2 dimers were predicted using the AlphaFold3 server (https://alphafoldserver.com/about)(76). As a reference for VP2 folding, the PDB of RVA VP2 strain RRV was used (6OGZ, https://doi.org/10.2210/pdb6OGZ/pdb).

### IDR predictions

The intrinsically disordered regions of proteins were determined with PONDR (Molecular Kinetics, Inc., https://www.pondr.com/) using the VSL2 algorithm. Data were plotted with GraphPad Prism [version 10.4.2].

### Immunofluorescence

MA/cytBirA and LMH cells were transfected and treated for immunofluorescence, as described previously by Lee et al., 2024 (65). For VLS formation composed of NSP5 and VP2, a ratio of 2:1 was used, with 2 µg and 1 µg of DNA plasmids, respectively. With the exception of VLS(VP2)i of RVC and RVI, which were obtained with a transfection ratio of NSP5 and VP2 of 1:4 and 1:2, respectively (**Fig. S2b and c**). VLS composed of NSP5, VP2, and NSP2 were obtained with a ratio of 2:1:1 using 2 µg, 1 µg, and 1 µg of DNA plasmids, respectively. The images were acquired using a confocal laser scanning microscope (DM550Q, Leica). Data were analyzed with Leica Application Suite (Mannheim, Germany) and ImageJ2 (version: 2.16.0/1.54p, https://imagej.net/software/imagej2/).

### Immunoblotting

Cell lysis and immunoblotting procedures were performed as described by Lee et al., 2024 (65).

### Detection of Halo-tagged proteins

MA104 cells seeded at a density of 2×10^5^ cells per well in a 12-multiwell plate. The cells were infected with vvT7.3 (MOI: 1 PFU/cell), followed by transfection with 1 µg DNA plasmid using 3 µl of Lipofectamine 2000 (Thermo Fisher Scientific) according to the manufacturer’s instructions. At 16 hpt, the cells were lysed in 30 µl TNN buffer (100 mM Tris-HCl, pH 8.0, 250 mM NaCl, 0.5% nonidet P-40, and cOmplete protease inhibitor cocktail (Roche, Switzerland) for 10 min on ice. The cell lysate was centrifuged at 17,000 x g for 7 min at 4°C. Then, 10 µl of supernatant was incubated with 10 µl of 2.5 µM HaloTag^®^TMR^™^Direct Ligand (Promega) in DMSO. The sample was incubated for 30 min in the dark at room temperature, followed by the addition of 10 µl of sample buffer (8% SDS, 40% glycerol, 200 mM Tris-HCl pH 6.8, 0.8% bromophenol blue, and 5 mM 2-mercaptoethanol). The samples were heated at 70°C for 3 min and migrated in an SDS-polyacrylamide gel followed by acquisition at 520 nm channel at Odyssey M Imager (LI-COR Biosciences).

### NanoBRET protein-protein interaction

HEK-293T cells, seeded at 8×10^5^ cells per well in 6-well plates. At 4 h post-seeding, the cells were transfected in a ratio HaloTag: NanoLuc of 10: 1, by adding 2000 ng and 200 ng of the respective DNA plasmids, using 6 µl of Lipofectamine LTX transfection reagent (ThermoFisher Scientific) diluted in 100 µl of Opti-MEM reduced medium. The transfection mixture was incubated for 30 min at room temperature and added to the cells. At 20 hpt, the cells were counted and diluted to 2×10^5^ cells per ml in 4% FCS in OptiMEM-I reduced serum medium. Then, 500 µl of diluted cells were mixed with 0.5 µl of 0.1 mM HaloTag^®^NanoBRET^™^618 Ligand (+ ligand, Promega) or 0.5 µl DMSO (-Ligand). Then, 40 µl of each mixture was distributed in quadruplicated in a white wall 384-wells plate. The cells were incubated for 6 h at 37°C and 5% CO2. Afterward, 10 µl of 5X solution of NanoBRET Nano-Glo substrate in Opti-MEM reduced serum medium was added per well. The luminescence was measured, in a range of 10 minutes, at 460 nm and 618 nm for donor emission and acceptor emission, respectively, using a Spark instrument (TECAN). The BRET ratio corresponds to the mean corrected mBU, which is obtained as follows:

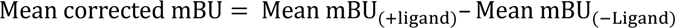

Where: *mBU* = (618 *nm*/460 *nm*) × 1000.

Statistical analysis was performed using:

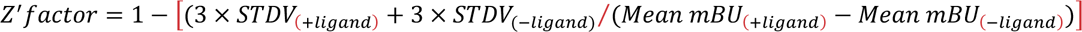

### Phylogenetic tree analysis

The CDS for rotavirus VP2 proteins were translated *in silico* into amino acid sequences using EMBOSS “transeq” (http://emboss.open-bio.org). The protein sequences were aligned using “mafft” [MAFFT v7.475 (23 November 2020); https://mafft.cbrc.jp/alignment/software/], and the aligned protein sequences were backtranslated using EMBOSS “transeq” (http://emboss.open-bio.org)(77, 78). The phylogeny from the nucleotide multiple sequence alignments was then inferred by using “BEAUTi” and “BEAST” (v1.10.4)(79). In brief, 10,000,000 Markov chain Monte Carlo (MCMC) steps with the Juke-Cantor model were performed, saving each 10,000th tree. After the burn-in of 100,000 states, the consensus tree for VP2 was calculated and visualized using FigTree (https://beast.community).

## Acknowledgments

This work was supported by the Swiss National Science Foundation grant # 10.005.230 to C. E. The funders had no role in study design, data collection, interpretation, or the decision to submit the work for publication.

## Author contributions

The authors declare no conflict of interest.

Conceptualization: ML, AC, CE; Methodology: ML, AC, CE; Software: KT, CA; Validation: ML, AC, KT, CA, CE; Formal analysis:AC, ML, KT, CA, CF, CE; Investigation: ML, AC, KT, CA, CE; Resources: CE; Data Curation: CE; Writing-Original Draft: CE; Writing-Review & Editing: ML, AC, KT, CA, CF, CE; Visualization: CE; Supervision: CE; Project Administration: CE; Funding Acquisition: CF, CE.

